# A modular architecture of human cellular aging across organs and diseases

**DOI:** 10.64898/2026.08.12.744367

**Authors:** Jing Guo, Cong-Cong Liu, Xicheng Yang, Jiahao Feng, Jia-Hao Wang, Wei Shi, Xian-lin Yu, Danqing Huang, Shan-Shan Dong, Yan Guo, Tie-Lin Yang

## Abstract

Aging is a heterogeneous biological process in which different cellular systems undergo molecular remodeling at distinct rates, yet whether human cellular aging follows an organized architecture across organs remains unclear. Here, we integrate a multi-organ human single-cell transcriptomic atlas with plasma proteomic profiles from approximately 50,000 participants to reconstruct cellular aging states at population scale. By projecting cell-type-enriched molecular signatures onto circulating proteins, we characterize aging patterns across 128 organ-cell type pairs and identify 14 cellular aging modules comprising conserved cross-organ programs and organ-specific aging states. These modules reveal cellular identity as a dominant organizing axis of human aging that transcends anatomical boundaries. Module-level aging states uncover substantial inter-individual heterogeneity, with 34% of individuals exhibiting extreme aging deviation in at least one cellular module. Cellular aging modules exhibit distinct temporal trajectories, with structural and tissue-resident modules showing earlier remodeling than immune lineages. The modular organization of cellular aging is reflected in disease susceptibility, with accelerated aging of specific modules, particularly epithelial aging, showing broad associations with disease burden and mortality. Longitudinal analyses further demonstrate the stability and clinical relevance of cellular aging states, whereas lifestyle, metabolic and pharmacological factors show selective relationships with individual aging programs. Together, our study establishes a modular framework for understanding human cellular aging and reveals an organization of biological aging that may help explain individual differences in healthspan.

## Introduction

Aging is the major risk factor for a wide range of chronic diseases and mortality^1^, yet chronological age provides only a limited description of biological decline. Individuals of the same chronological age can exhibit striking differences in physiological function, molecular states and disease susceptibility, indicating that aging proceeds along heterogeneous trajectories rather than a uniform path^2^. This heterogeneity has motivated the development of biological aging measures based on molecular, physiological and clinical features, which have revealed substantial variation between individuals in the pace of aging and vulnerability to age-related diseases^3^.

Recent advances in aging clocks based on DNA methylation, transcriptomics, proteomics, metabolomics and imaging have demonstrated that biological aging can be quantified beyond chronological age, which have substantially improved our ability to estimate individual aging trajectories and predict health outcomes^4–9^. However, most existing measures summarize aging into a single composite score or organ-level estimate, thereby obscuring the cellular processes that generate biological heterogeneity. Because tissues are composed of diverse cell populations with distinct functions, regulatory programs and stress responses, understanding aging at cellular resolution is essential for revealing the biological basis of individual differences in aging^10,11^.

Single-cell studies in model organisms and human tissues have begun to reveal that aging is accompanied by cell-type-specific molecular remodeling, including alterations in cellular composition, inflammatory states, regenerative capacity and tissue maintenance programs^12–18^. These studies suggest that aging is not synchronized across all cellular populations; instead, different cell types may possess distinct vulnerabilities and trajectories across the lifespan^12,19–22^. However, an important unanswered question remains: do cellular aging processes occur as independent changes restricted to individual cell populations, or are they organized into conserved biological programs that coordinate aging across organs?

Addressing this question in humans has been challenging because direct measurement of cellular aging requires access to tissues, limiting large-scale longitudinal investigation. Circulating plasma proteins provide an accessible window to overcome this limitation because they integrate molecular signals originating from diverse cellular systems and can be measured in large population cohorts^23–26^. Recent studies have demonstrated that plasma proteomic profiles can capture cell-type-associated aging signatures and provide non-invasive estimates of cellular biological age^26^. However, whether these heterogeneous cellular aging signals form higher-order organizational structures across tissues, and how such structures relate to health trajectories, remains unclear.

Here, we integrate a multi-organ human single-cell transcriptomic atlas from Tabula Sapiens with large-scale plasma proteomics profiles from ∼50,000 UK Biobank participants to reconstruct the organization of human cellular aging at population scale^27–29^. We develop aging state models across organ-cell type pairs and characterize their relationships to identify cellular aging modules. We reveal a modular architecture of human cellular aging consisting of conserved cross-organ programs and organ-specific aging states. We further show that cellular aging modules exhibit distinct temporal trajectories and inter-individual heterogeneity, with module-specific aging patterns associated with disease burden, incident disease, mortality risk, and modifiable health-related factors. Together, our findings provide a framework for understanding how cellular aging is organized across the human body and why individuals exhibit divergent aging trajectories.

## Results

### Human cellular aging is organized into conserved cross-organ modules

To determine whether human cellular aging follows an organized architecture across tissues, we developed a population-scale framework integrating multi-organ single-cell transcriptomic references with plasma proteomic profiles (Figure 1). Rather than focusing on individual aging predictors, we sought to characterize how cellular aging trajectories are coordinated across different organ and cell-type contexts. Briefly, we (i) integrated multi-organ single-cell transcriptomic and plasma proteomic data to define organ-cell type-enriched protein features and derive organ-cell type-level age gaps using age-prediction models, where the age gap represents an individual’s relative biological age compared with age-matched peers; (ii) identified cellular aging modules based on the correlation structure of organ-cell type-level age-gaps and developed module-level aging state model; and (iii) evaluated associations of these modules with disease outcomes, mortality risk, and health-related phenotypes. See Tables S1-S8 for detailed descriptions of the single-cell transcriptomic, plasma proteomic, age-associated protein, and clinical and phenotypic datasets used in the study.

**Figure 1.**
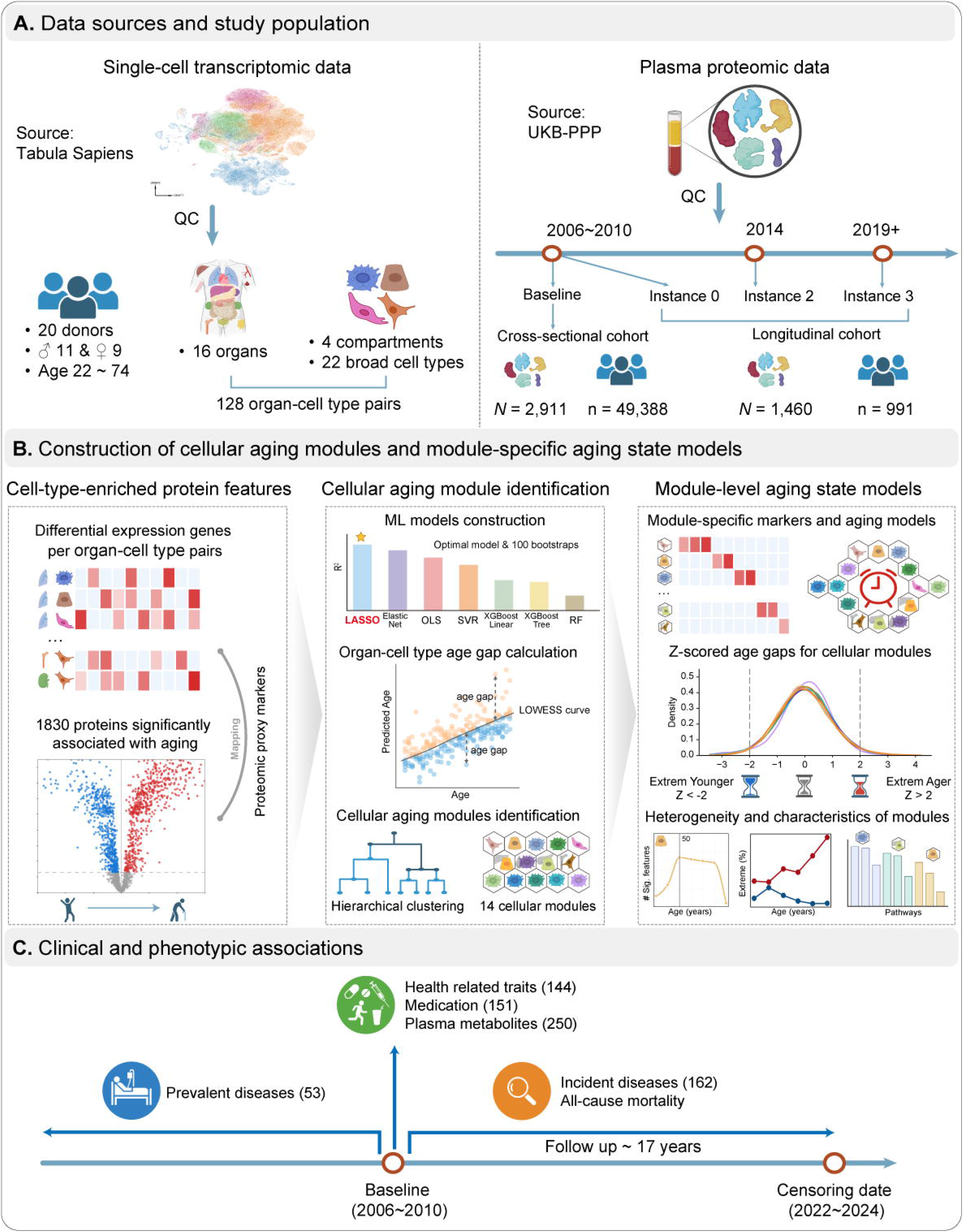
Overview of the study workflow. (A) Data sources and study population. Multi-organ single-cell transcriptomic data from Tabula Sapiens were integrated with UK Biobank plasma proteomic data from cross-sectional (n = 49,388) and longitudinal (n = 991) cohorts. (B) Construction of cellular aging modules and module-specific aging states. Cell-type-enriched signatures from single-cell transcriptomics were mapped to plasma proteins to construct organ–cell type-specific aging models. Hierarchical clustering of organ-cell type age gaps identified 14 cellular aging modules. Module-specific protein features were then used to estimate module-level aging states. Standardized age gaps were used to define extreme aging states (z-score ≥ 2 for accelerated aging and z-score ≤ −2 for decelerated aging). These modules were further characterized by their population heterogeneity and biological functions. (C) Association of cellular aging modules with health and disease. Module-specific age gaps were assessed for associations with diseases, mortality, health-related traits, medications, and circulating metabolites. Schematics were created with BioRender.com.

After quality control, the single-cell transcriptomic datasets comprised 16 human organs and 22 broad cell types, yielding 128 organ-cell type pairs spanning diverse cellular compartments (Figure S1). To define organ-cell type-enriched proteomic features (Table S9), we identified differentially expressed genes for each organ-cell type pair relative to all other organ-cell types and projected these signatures onto a subset of plasma proteins associated with chronological age, as determined by proteome-wide linear regression analyses (see STAR Methods). We estimated organ-cell type aging states and quantified individual deviations from age-matched expectations. Comparing seven machine learning algorithms for age prediction and selected LASSO regression based on overall performance (Table S10). Models were trained using bootstrap-aggregated LASSO with 100 resampling iterations and tested on held-out test samples. Among the 128 organ-cell type pairs, 107 aging models showed robust performance, with Pearson correlation coefficients (*r*) ranging from 0.22 to 0.85 and mean absolute errors (MAE) ranging from 3.3 to 6.5 years (Figure S2 and Table S11). These models provided a quantitative representation of cellular aging variation across individuals and enabled investigation of whether aging trajectories were independently distributed or coordinated across cellular systems.

We next asked whether cellular aging trajectories represented isolated changes restricted to individual cell populations or whether they converge into conserved biological programs. Hierarchical clustering of organ–cell type aging states revealed 14 distinct cellular aging modules, including 10 cross-organ modules and four organ-specific modules (Figures 2A and S3). Cross-organ modules were primarily composed of shared cellular lineages, including epithelial cells, endothelial cells, stromal cells, and multiple immune lineages, whereas organ-specific modules comprised specialized cellular populations, including hepatocytes, pancreatic epithelial cells, hematopoietic cells and bone marrow stem cells.

**Figure 2.**
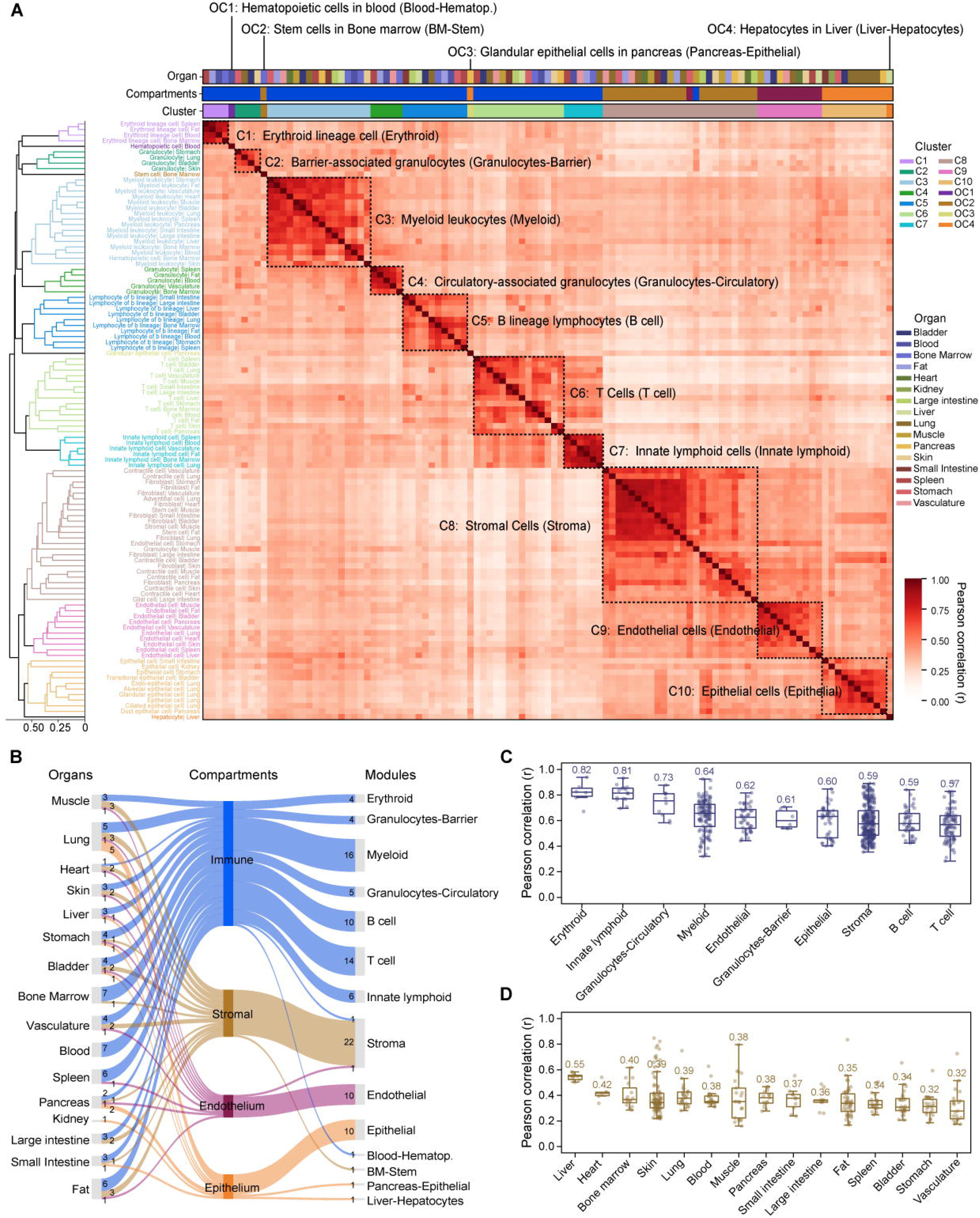
Identification of cross-organ and organ-specific cellular aging modules. (A) Hierarchical clustering of organ–cell type aging states based on age-gap similarity. Left, dendrogram showing hierarchical clustering of organ–cell type pairs using pairwise Pearson correlations of age gaps across individuals. Right, heatmap displaying Pearson correlation coefficients among organ-cell type-level age gaps, ordered according to clustering results. Dashed boxes indicate identified cellular aging modules. Colored annotations indicate organ origin, cellular compartment, and module assignment. (B) Distribution of organ–cell type pairs across organs, cellular compartments, and cellular aging modules. Sankey diagram showing the distribution of organ-cell type pairs to cellular aging modules. (C) Within-module similarity of organ-cell type aging states. Box plots show the distribution of pairwise Pearson correlation coefficients among organ-cell type age gaps within each cellular aging module. Mean correlation coefficients (ì) are indicated above each module. (D) Within-organ similarity of organ-cell type aging models. Box plots show the distribution of pairwise Pearson correlation coefficients among organ-cell type age gaps within each organ. Mean correlation coefficients (μ) are indicated above each organ. Module abbreviations: Erythroid lineage cell (Erythroid), barrier-associated granulocytes (Granulocytes-Barrier), myeloid leukocytes (Myeloid), circulatory-associated granulocytes (Granulocytes-Circulatory), B lineage lymphocytes (B cell), T Cells (T cell), innate lymphoid cells (Innate lymphoid), stromal Cells (Stroma), endothelial cells (Endothelial), epithelial cells (Epithelial), blood hematopoietic cells (Blood-Hematop.), bone marrow stem cells (BM-Stem), glandular epithelial cells in pancreas (Pancreas-Epithelial), and hepatocytes in liver (Liver-Hepatocytes).

Strikingly, cellular aging patterns were organized predominantly according to cellular identity rather than anatomical origin. Cell populations sharing similar developmental and functional programs clustered together despite residing in different organs, indicating that cell identity represents a major organizing axis of cellular aging, despite differences in tissue context. For example, epithelial populations from lung, kidney, and gastrointestinal tract converged into a shared epithelial aging module, whereas immune populations across blood, spleen, bone marrow and multiple tissues formed conserved immune aging programs.

Although cellular identity represented the primary organizing principle, specialized organ functions introduced additional layers of aging heterogeneity. Organ-specific modules captured aging patterns of highly specialized cellular populations, such as hepatocytes involved in metabolic and detoxification functions, pancreatic epithelial cells responsible endocrine and exocrine secretory regulation, and bone marrow stem cells involved in hematopoietic maintenance. These findings suggest a hierarchical architecture of cellular aging, in which conserved lineage-associated programs provide the fundamental framework while tissue-specific physiological environments further shape individual aging trajectories.

Modules showed biologically interpretability by largely following cell identity, while rare exceptions highlighted aging programs that extend beyond conventional cell-type annotations (Figure 2B). For example, granulocytes in muscle and endothelial cells in the stomach clustered within the stromal module. This convergence may reflect shared biological processes involving tissue remodeling, inflammatory regulation, and extracellular matrix interactions^30,31^. In addition, gene set enrichment analysis confirmed that the identified cellular aging modules retained distinct biological identities (Figure S4).

Furthermore, age-gap correlations were substantially higher among organ-cell type pairs assigned to the same module than among pairs grouped by organ origin (mean within-module correlation *r* = 0.59-0.82 versus organ-based correlation *r* = 0.32-0.55, *p*-value = 5.81 ×·10^-104^, Mann-Whitney U test) (Figures 2C and 2D), supporting cellular identity as a strong determinant of aging patterns than tissue location.

Together, these findings reveal that human cellular aging is not a collection of independent cellular alterations but a structured biological system composed of conserved cross-organ and organ-specific aging programs.

### Cellular aging modules undergo asynchronous remodeling across adulthood

We next sought to characterize how these cellular aging modules change across adulthood and whether different cellular systems follow distinct aging trajectories. To quantify biological variation at the module level, we developed module-specific aging state models. For each of the 14 cellular modules, module-specific protein features were used to train independent LASSO bootstrap models for chronological age prediction and derive module-level age gaps (Figure 3A and S5; Table S12). Predictive performance in the test set ranged from moderate to strong (Pearson’s *r* = 0.23-0.83; MAE = 3.48-6.51 years). Comparable performance was observed using alternative machine-learning algorithms (Table S13), supporting the robustness of module-level aging state estimation. To evaluate model reproducibility over time, we trained module-level aging state using a reduced Olink panel comprising 1,459 proteins (see STAR Methods). Despite the reduced feature set, predictive performance remained comparable to the full proteomic models (Pearson’s *r* = 0.22-0.79; MAE = 4.04-6.72 years; Table S14). In an independent longitudinal cohort with repeated proteomic measurements, module-specific age gaps exhibited moderate-to-high stability over approximately 15 years (*r* = 0.28-0.70; mean *r* = 0.56; Figure S6), indicating that cellular aging modules represent relatively stable individual characteristics rather than transient molecular fluctuations.

**Figure 3.**
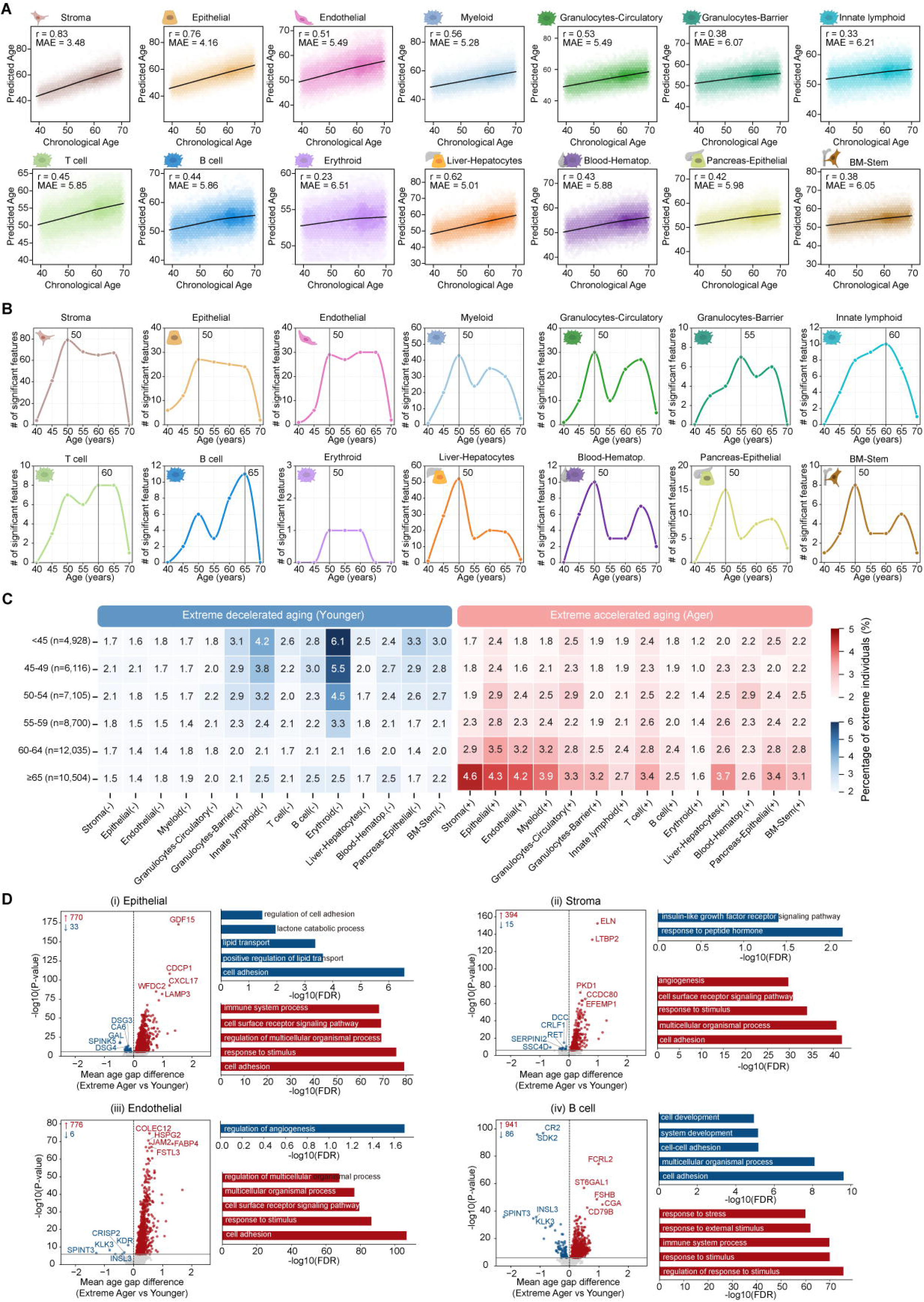
Module-specific aging states reveal temporal dynamics and heterogeneity of cellular aging. (A) Performance of module-specific aging state models. Independent aging models were constructed for each cellular aging module using module-specific age-associated plasma proteins. Hexbin plots show predicted age versus chronological age in the test datasets, with locally weighted scatterplot smoothing (LOWESS) curves overlaid. Pearson correlation coefficients (*r*) and mean absolute errors (MAE) are indicated for each model. (B) Temporal dynamics of molecular aging across cellular aging modules. Differential expression-sliding window analysis (DE-SWAN) was performed using module-specific protein features across chronological age. Curves show the number of significantly age-associated proteins across sliding-window center ages. Vertical dashed lines indicate peak of molecular remodeling. (C) Age-dependent pattern of extreme cellular aging states. Heatmaps show the proportions of accelerated agers (red; age-gap z-score ≥ 2) and decelerated agers (blue; age-gap z-score ≤ −2) across six chronological age groups in the UK Biobank cross-sectional cohort (n = 49,388) for each of the 14 cellular aging modules. (D) Functional features associated with extreme cellular aging in representative modules. Volcano plots show proteins differentially expressed between accelerated and decelerated agers in epithelial (i), stromal (ii), endothelial (iii), and B-cell (iv) modules. Differential expression was assessed using Bonferroni-adjusted significance threshold (*P* < 1.23 × 10^-6^). Top five significant proteins are labeled, and the numbers of significantly accelerated (red) and decelerated (blue) agers are indicated. Right, Gene Ontology Biological Process (GOBP) enrichment analysis of differentially proteins, showing the top five enriched biological processes. Red and blue indicate enrichment in accelerated and decelerated agers, respectively.

We next investigated whether different cellular aging modules undergo molecular remodeling at similar stages of adulthood. Pairwise correlations among module-specific age gaps were generally positive but moderate (*r* = 0.18-0.55; Figure S7), indicating that aging processes across cellular systems are partially coordinated across cellular modules while retaining substantial module-specific dynamics.

To determine when distinct cellular aging programs undergo their most pronounced molecular remodeling, we performed differential expression-sliding window analysis (DE-SWAN) using module-specific protein features. Distinct temporal patterns emerged across cellular aging modules (Figure 3B). Structural and tissue-resident modules, including stromal, epithelial, endothelial, and organ-specific parenchymal cells, exhibited the earliest remodeling peaks at approximately 50 years of age. In contrast, immune-related modules displayed progressively delayed peaks, from myeloid and granulocytes-circulating (∼50 years), to granulocyte-barrier (∼55 years), innate lymphoid cells (∼60 years), and finally adaptive immune compartments, with T cell and B cell modules peaking at approximately 60 and 65 years, respectively. These results indicate that cellular aging does not proceed as a synchronized systemic process, but instead follows a temporal hierarchy in which different cellular systems enter aging-associated states at distinct stages of life.

To characterize how these temporal differences translate into individual variation, we examined the distribution of extreme aging states across the population. Individuals were stratified into extreme accelerated (z-score ≥ 2) and decelerated (z-score ≤ −2) agers based on module-specific age-gap z-scores, revealing substantial heterogeneity in cellular aging trajectories (Figure 3C). Across the 14 modules, the two extreme groups showed a mean age-gap separation of approximately 17.9 years, with the largest difference observed in the epithelial module (22.1 years; Figure S8). Approximately 34% of individuals exhibited extreme aging deviation in at least one module, including 20.8% with isolated acceleration in a single-module and 13.2% displaying coordinated acceleration across multiple modules (Figure S8). A small subset of individuals simultaneously displayed accelerated aging in some modules and decelerated aging in others, indicating marked within-individual heterogeneity across cellular systems.

The prevalence of extreme aging states also differed substantially across modules and chronological age groups. Rather than following a uniform trajectory, cellular modules exhibited four distinct aging patterns (Figure S9). For example, epithelial, endothelial, and stromal modules showed a gradual accumulation pattern with progressively increasing accelerated agers across adulthood. Barrier granulocyte, bone marrow stem cell, and pancreatic epithelial modules exhibited late-life acceleration, characterized by a marked increase in accelerated agers after approximately 60 years of age. The B cell module displayed dynamic fluctuations throughout adulthood, whereas the erythroid module showed a young-state loss dominant pattern, characterized by a progressive decline in decelerated agers with only modest increases in accelerated agers.

To understand the molecular basis underlying these distinct aging trajectories, we compared plasma protein signatures between individuals with extreme accelerated and decelerated aging states for each module (Figure 3D). Representative modules exhibited distinct functional remodeling between the two aging states. Accelerated epithelial aging was characterized by a transition from lipid homeostasis-related processes toward inflammatory and immune pathways, whereas the endothelial module shifted from angiogenesis and vascular development toward cell adhesion and stimulus-responsive signaling. Similarly, the stromal module shifted from endocrine and metabolic signaling toward tissue organization and cell adhesion-related processes, whereas the B cell module transitioned from developmental and homeostatic processes toward immune system and immune response pathways.

Together, these findings demonstrate that human cellular aging is a temporally structured and heterogeneous process. Different cellular systems undergo molecular remodeling at distinct stages of adulthood, generating individualized combinations of aging states that contribute to variation in biological aging trajectories.

### Cellular aging modules are associated with disease burden and progression

We next investigated whether the modular organization of cellular aging was reflected in human disease patterns. We evaluated associations between module-specific age gaps and 53 prevalent diseases in the UK Biobank cohort. Overall, 545 of 742 associations (73.5%) reached statistical significance, indicating widespread relationships between cellular aging and diseases (Figure 4A and Table S15). However, the magnitude and spectrum of these associations varied considerably across modules. Among the 14 cellular aging modules, the epithelial module exhibited the broadest disease associations (51/53 diseases), particularly for metabolic and cardiovascular disorders, including type 2 diabetes (β = 1.02, *p*-value = 5.72 × 10⁻^245^) and hypertension (β = 0.46, *p*-value = 1.81 × 10^-174^) (Figure 4B). The broad biological relevance of epithelial aging is consistent with the central role of epithelial tissues in maintaining barrier integrity, regulating tissue repair and mediating interactions between organisms and external environment. These findings suggest that epithelial aging may represent a shared biological interface linking cellular aging processes with diverse chronic disease states. Other cellular aging modules displayed more selective disease association patterns. For example, the myeloid module showed stronger relationships with inflammatory and immune-related diseases such as asthma (β = 0.33, *p*-value = 3.31 × 10⁻^48^), whereas B cell module was associated relatively few diseases (Figure 4B).

**Figure 4.**
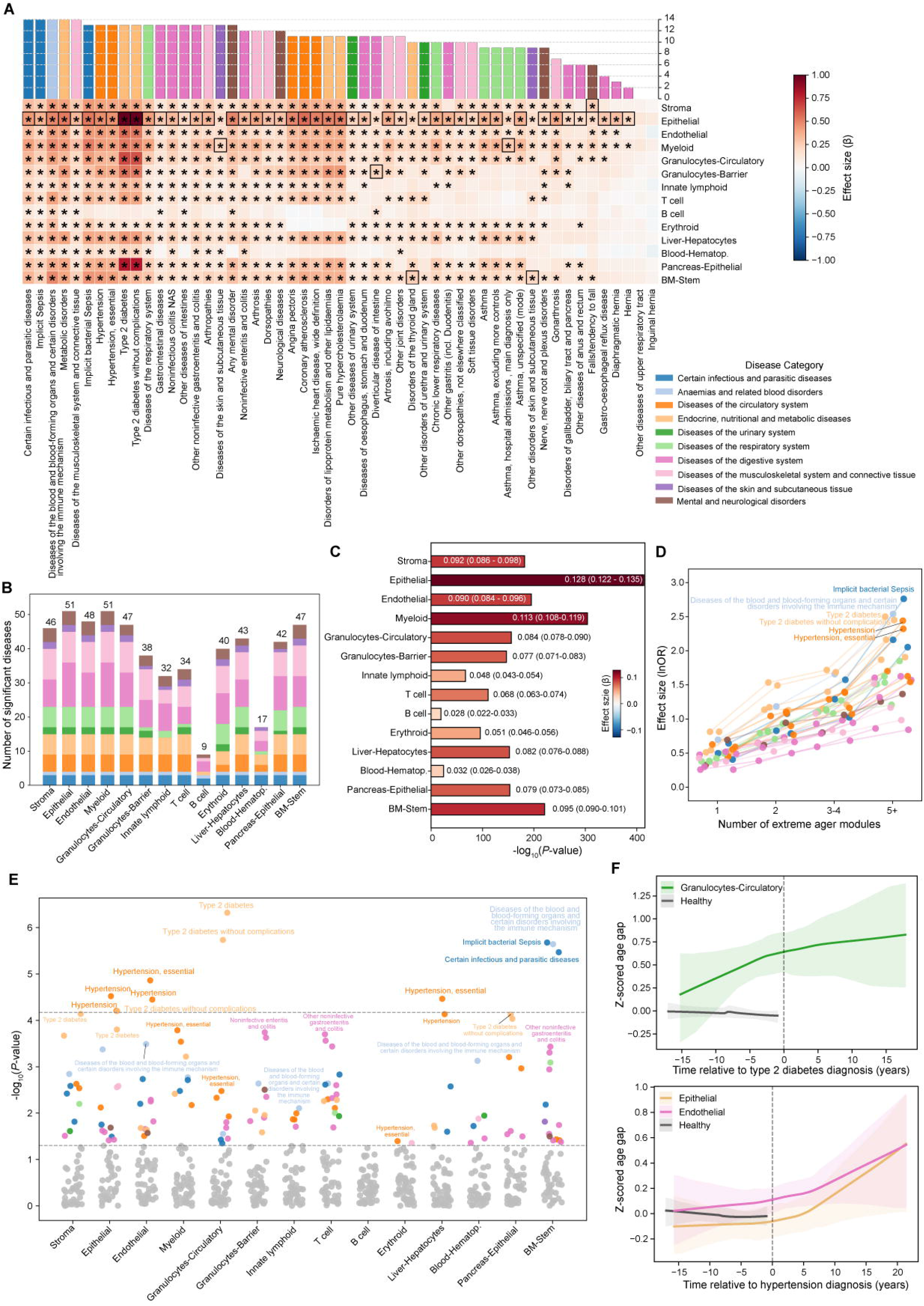
Cellular aging modules link biological aging to disease burden and progression. (A) Associations between prevalent diseases and module-specific age gaps. Heatmap showing associations between prevalent diseases and the 14 cellular aging modules estimated using linear regression models adjusted for covariates. Colors represent regression coefficients (β). Asterisks denote Bonferroni-corrected significant associations (*P* < 4.69 × 10^-6^; 0.05 divided by 14 modules and 761 phenotypes). Black borders highlight the module showing the strongest association for each disease. The upper bar plot indicates the number of modules significantly associated with each disease. (B) Stacked bar plot showing the number of significantly associated prevalent diseases for each cellular aging module, stratified by disease category. (C) Association between multimorbidity burden and cellular aging modules. Bar plot showing associations between the number of prevalent disease categories and module-specific age gaps estimated using linear regression models. The x-axis represents -log_10_(*p*-value), and colors indicate regression coefficients (β). Regression coefficients and 95% confidence intervals (CIs) are shown for each module. (D) Associations between prevalent diseases and extreme modular aging burden. Scatter plot showing disease associations with the number of extremely aged modules estimated using logistic regression models adjusted for covariates. The x-axis represents the number of extremely aged modules, and the y-axis shows effect sizes. Lines connect the same disease across different aging burden levels. Colors indicate disease categories. The six diseases with the largest effect sizes among individuals with ≥5 extremely aged modules are labeled. (E) Associations between disease progression and module-specific age gaps. Associations between years since diagnosis and module-specific age gaps were assessed among individuals diagnosed before proteomic sampling using linear regression models adjusted for covariates. The x-axis represents cellular aging modules and the y-axis represents -log_10_(*p*-value). Significant disease – module associations after Bonferroni correction are labeled, with the strongest nominal association for each module additionally annotated. Gray points indicate non-significant associations. Dashed lines represent Bonferroni-corrected (*P* < 6.74 × 10^-5^; 0.05 divided by 14 modules and 53 prevalent diseases) and nominal significance thresholds. (F) Longitudinal dynamics of cellular aging surrounding disease diagnosis. LOWESS curves show changes in module-specific age gaps relative to disease diagnosis, with shaded regions indicating 95% confidence intervals estimated by bootstrap resampling. Colored curves represent disease-associated cellular aging modules, whereas gray curves represent controls. The x-axis represents time relative to disease diagnosis, with negative and positive values representing pre- and post-diagnostic periods, respectively. Upper panels show type 2 diabetes-associated changes in the Granulocytes-Circulatory module; lower panels show hypertension-associated changes in the Epithelial and Endothelial modules.

To further assess overall disease burden, we examined associations between multimorbidity and module-specific age gaps. Module-specific age gaps were positively associated with multimorbidity, defined as the number of diagnoses across 11 disease categories (Figure 4C). The epithelial module showed the strongest association (β = 0.128, *p*-value < 1 × 10⁻^400^), whereas the B cell module showed the weakest (β = 0.028, *p*-value = 5.07 × 10⁻^20^). Similar results were obtained using proportional odds logistic regression after categorizing multimorbidity into five levels (Figure S12). In analyses, cellular age gaps increased progressively with increasing disease burden, with stronger associations observed among younger participants (40-49 years).

We next investigated whether extreme cellular aging identified individuals with greater disease burden. Logistic regression analyses of both single-module and multi-module extreme aging states identified 204 significant associations among 1,702 tests (Figures 4D, S10, and S11; Tables S16 and S17). Among individuals with single-module extreme aging, type 2 diabetes and hypertension showed associations with extreme aging in several distinct cellular modules, including epithelial, myeloid, granulocyte, stromal, and pancreas-specific modules. In individuals with multi-module extreme aging, association strengths increased progressively with the number of extremely aged modules, particularly for type 2 diabetes, hypertension, implicit bacterial sepsis, and hematological disorders, indicating that disease burden accumulates with increasing cellular aging across multiple biological systems.

We next examined whether cellular aging was associated with disease progression by relating module-specific age gaps to disease duration among individuals diagnosed before blood sampling (Figure 4E and Table S18). Longer disease duration was associated with progressively higher age gaps in the epithelial, endothelial, granulocytes-circulatory, and liver-hepatocyte modules among individuals with type 2 diabetes or hypertension. In contrast, longer disease duration was associated with lower bone marrow stem cell age gaps in individuals with implicit bacterial sepsis, infectious diseases, and hematological disorders.

Finally, we characterized longitudinal changes in module-specific age gaps by aligning repeated proteomic measurements relative to disease diagnosis (Figure 4F). Distinct module-specific trajectories emerged across diseases. For example, the granulocyte-circulatory module exhibited sustained increases throughout progression of type 2 diabetes, whereas epithelial and endothelial age gaps increased progressively around and after hypertension diagnosis. Additional examples are shown in Figure S13, illustrating diverse temporal patterns across diseases and cellular modules. Together, these findings demonstrate that disease-associated cellular aging follows diverse module-specific trajectories, with some aging signatures preceding clinical diagnosis and others continuing to accumulate after disease onset.

### Module-specific cellular aging predicts incident disease and all-cause mortality

We next evaluated whether module-specific cellular aging could predict future disease incidence using Cox proportional hazards models. Across 162 incident diseases, 1,668 of 2,268 associations (73.5%) were significantly positive, indicating that cellular aging states capture broad patterns of future disease vulnerability (Figure 5A and Table S19). Among all modules, the epithelial module exhibited the broadest disease associations (151/162 diseases), with particularly strong effects for type 2 diabetes (hazard ratio (HR) = 1.49, *p*-value = 4.48 × 10⁻^138^), and chronic obstructive pulmonary disease (HR = 1.64, *p*-value = 1.45 × 10⁻^152^). Other modules displayed preferential associations with distinct disease categories, including the stromal module with cardiovascular diseases, the myeloid module with inflammatory and infection-related diseases, and the bone marrow stem cell module with cerebrovascular diseases. Incident disease associations were broadly distributed across cellular aging modules, although both the strength and spectrum of associations varied substantially among modules (Figure 5B).

**Figure 5.**
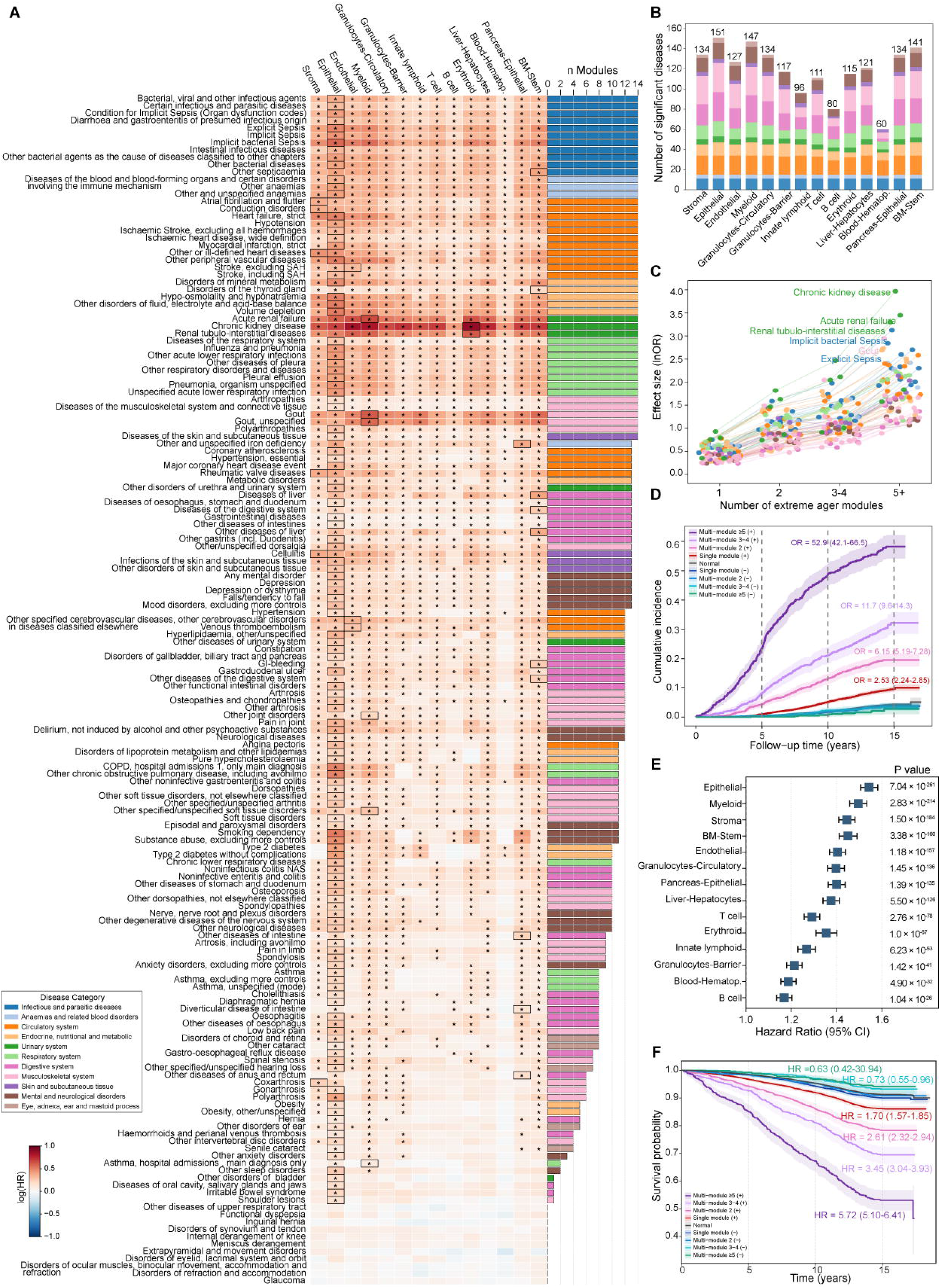
Cellular aging modules are associated with incident disease and mortality risk. (A) Associations between module-specific age gaps and incident diseases. Heatmap showing associations between cellular aging modules and incident diseases estimated using Cox proportional hazards regression models. Colors represent log-transformed hazard ratios (log HR). Asterisks denote Bonferroni-corrected significant associations (*P* < 4.69 × 10^-6^; 0.05 divided by 14 modules and 761 diseases). Black borders highlight the most strongly associated module for each disease. The right-side bar plot indicates the number of modules significantly associated with each disease. (B) Stacked bar plot showing the number of significantly associated incident diseases for each cellular aging module, stratified by disease category. (C) Associations between extreme modular aging burden and incident diseases. Scatter plot showing associations between the number of extremely aged modules and incident diseases estimated using logistic regression models adjusted for covariates. The x-axis indicates the number of extremely aged modules, and the y-axis represents effect sizes. Lines connect the same disease across different aging burden levels. Colors indicate disease categories. The six diseases with the largest effect sizes among individuals with ≥5 extremely aged modules are labeled. (D) Cumulative incidence of chronic kidney disease according to multi-module extreme aging burden. Cumulative incidence function (CIF) curves for chronic kidney disease stratified by the number of extremely aged or extremely young cellular modules. The x-axis represents follow-up time, and the y-axis represents cumulative incidence. Groups were defined according to the number of extreme modules, including single-module, multi-module (2, 3-4, and ≥5 modules), and normal aging groups. Normal individuals had module age-gap z scores between -2 and 2 across all modules. *P* values were calculated using Gray’s test for equality of cumulative incidence functions. (E) Associations between module-specific age gaps and all-cause mortality. Forest plot showing hazard ratios (HRs) and 95% confidence intervals (CIs) for associations between the 14 cellular aging modules and all-cause mortality estimated using Cox proportional hazards regression models. Statistical significance was assessed using Bonferroni correction (*P* < 4.69 × 10^-6^; 0.05 divided by 14 modules and 761 diseases). (F) Survival according to multi-module extreme aging burden. Kaplan-Meier curves show survival probabilities according to the number of extremely aged cellular modules. Groups included single-module, multi-module (2, 3-4, and ≥5 modules), and normal aging groups. Hazard ratios indicate mortality risk in each extreme aging group compared with the normal aging group.

We next investigated whether extreme cellular aging identifies individuals at particularly high disease risk. Extreme aging within a single module yielded 171 significant associations among 3,623 tests (Figure S14 and Table S20), with the epithelial module accounting for nearly half of all significant associations (80/171). Individuals with extreme epithelial aging exhibited markedly increased risks of type 2 diabetes (odds ratio (OR) = 3.58, *p*-value = 1.50 × 10⁻^23^), chronic obstructive pulmonary disease (OR = 4.49, *p*-value = 1.96 × 10⁻^31^), and hypertension (OR = 2.12, *p*-value = 2.73 × 10⁻^13^). Cumulative incidence analyses further confirmed these findings, showing substantially higher 10- and 15-year disease incidence among individuals with extreme epithelial aging, whereas individuals with extremely young epithelial profiles consistently exhibited lower disease incidence (Figure S15 and Table S21). For example, the 15-year cumulative incidence of type 2 diabetes increased progressively from 2.7% in individuals with extremely young epithelial profiles to 4.2% in normally aging individuals and 18.3% in those with extreme epithelial aging.

We further examined disease risk increased with the accumulation of extreme aging across multiple cellular modules. Individuals with multiple extremely aged modules exhibited significant associations with 55.0% of incident diseases (89/162) (Figure S16 and Table S22). For most diseases, effect sizes increased progressively with the number of extremely aged modules, demonstrating a dose-dependent relationship between cumulative cellular aging burden and future disease risk (Figure 5C). Consistently, cumulative incidence analyses showed progressively increasing disease incidence with increasing numbers of extremely aged modules. For example, the 15-year cumulative incidence of chronic kidney disease increased from 4.3% in normally aging individuals to 9.7%, 19.5%, and 58.2% among individuals with one, multiple, and ≥5 extremely aged modules, respectively (Figure 5D and Table S23).

Finally, we evaluated associations between cellular aging and all-cause mortality. Age gaps in all 14 modules were positively associated with mortality risk, with the strongest effects observed for the epithelial (HR = 1.54, *p*-value = 7.04 × 10⁻^261^), myeloid (HR = 1.50, *p*-value = 2.83 × 10⁻^214^), and stromal modules (HR = 1.45, *p*-value = 1.50 × 10⁻^184^) (Figure 5E and Table S24). Extreme aging in several modules was likewise associated with increased mortality risk (Figure S17 and Table S25). Kaplan-Meier analyses further demonstrated progressively reduced survival with increasing cellular aging burden (Figure S18 and Table S26). Fifteen-year survival declined from 91.3% in normally aging individuals to 86.1%, 78.3%, and 53.1% among individuals with one, multiple, and ≥5 extremely aged modules, respectively, whereas individuals with extremely young modules consistently exhibited the highest survival probabilities (Figure 5F and Table S26). Together, these findings demonstrate that increasing cellular aging burden is associated with progressively higher risks of incident disease and all-cause mortality.

### Cellular aging modules are associated with modifiable health and metabolic factors

Finally, we investigated whether the modular organization of cellular aging was reflected in potentially modifiable health-related and metabolic factors. To reduce potential confounding by pre-existing disease, analyses were restricted to participants without the relevant diagnosed conditions at baseline. Across 144 lifestyle, physiological, socioeconomic, and behavioral traits (up to n = 18,712) we identified 167 significant associations with module-specific age gaps, comprising 104 positive and 63 negative associations (Figures 6A-F; Table S27).

**Figure 6.**
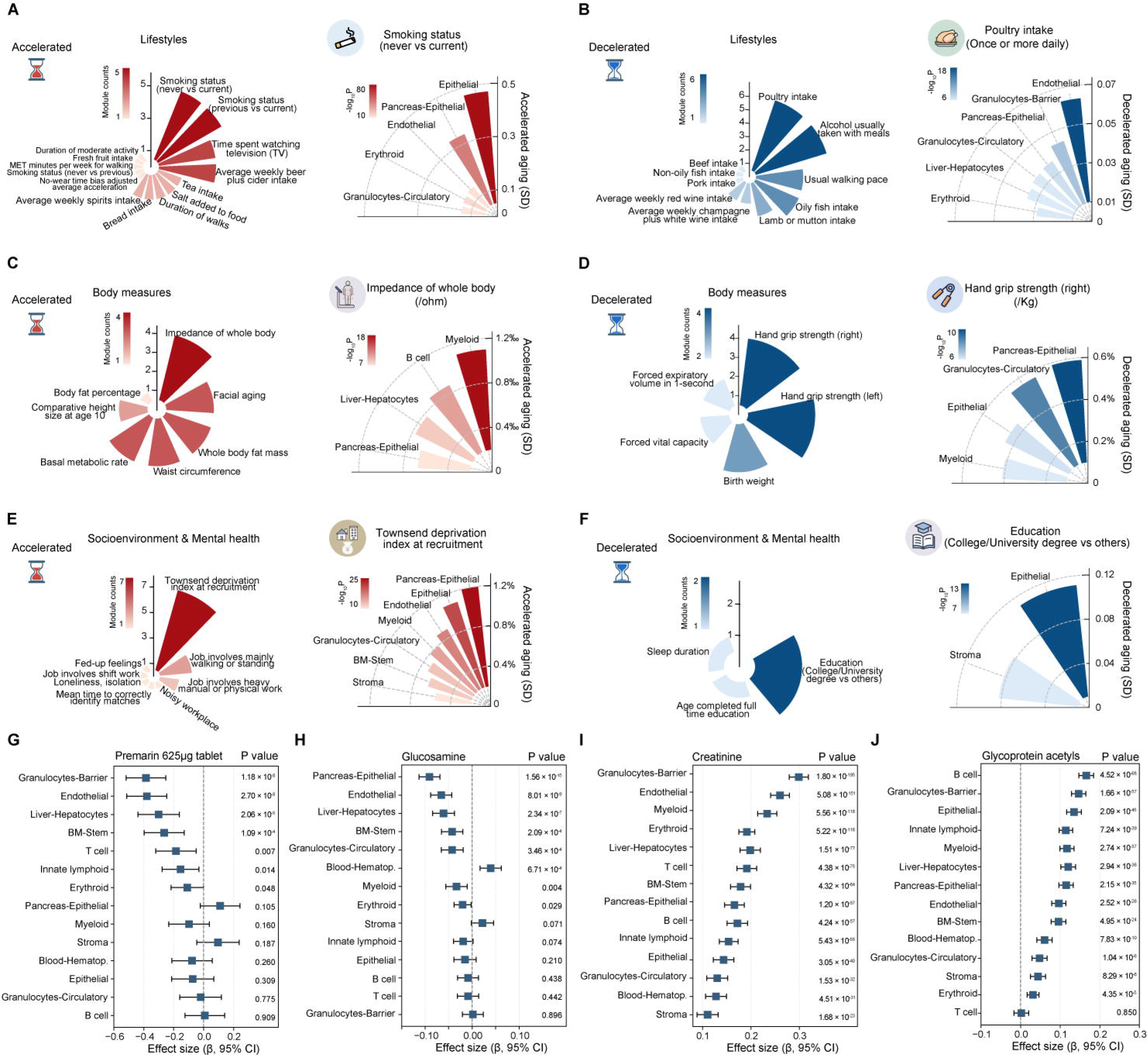
Associations of cellular aging modules with health-related traits, medications, supplements, and plasma metabolites. (A–F) Associations between cellular aging modules and health-related traits, including lifestyle (A, B), body measures (C, D), and socioenvironmental and mental health traits (E, F). Associations were estimated using covariate-adjusted linear regression models. The left plots show the number of cellular aging modules significantly associated with each trait, whereas the right plots display regression coefficients (β) for the trait associated with the largest number of modules. Red and blue indicate positive and negative associations, respectively. To facilitate visualization, negative regression coefficients are displayed as absolute β values. (G–J) Representative associations of medications and circulating metabolites with module-specific age gaps. Forest plots show regression coefficients (β), 95% confidence intervals (CIs), and *P* values for Premarin 625 μg tablet use (G), glucosamine use (H), creatinine (I), and glycoprotein acetyls (J). Statistical significance was assessed using Bonferroni correction (*P* < 4.69 × 10^-6^; 0.05 divided by 14 modules and 761 traits).

Several adverse lifestyle, physiological and socioeconomic factors were consistently associated with higher age gaps across multiple cellular aging modules. Current smoking was among the most broadly associated exposures and was associated with accelerating epithelial aging compared with never smoking (β = 0.470, *p*-value = 4.35 × 10⁻^91^). Higher whole-body impedance and greater socioeconomic deprivation were likewise associated with increased age gaps in multiple modules. In contrast, traits generally associated with healthy aging, including greater hand-grip strength, higher educational attainment, and healthier dietary habits, were associated with lower age gaps across multiple cellular systems. For example, daily poultry intake was associated with a lower endothelial age gap (β = -0.063, *p*-value = 1.91 × 10⁻^18^). Likewise, stronger hand-grip strength was associated with a lower pancreas-epithelial age gap, whereas individuals with a college or university degree exhibited younger epithelial biological age than those with lower educational attainment.

Not all associations were shared across modules. Several anthropometric, cardiovascular, and dietary traits displayed module-specific association patterns, with opposite directions observed across different cellular systems (Figure S19). For example, body mass index was negatively associated with the pancreas-epithelial age gap but positively associated with endothelial aging, whereas both systolic and diastolic blood pressure were associated with lower T cell age gaps but higher age gaps in endothelial and granulocyte-barrier modules. These findings indicate that health-related factors exert both common and module-specific influences on cellular aging.

Medication and supplement use also showed extensive associations with module-specific aging states. After multiple-testing correction, 545 significant associations (25.8%) were remained, involving 71 medications or supplements associated with higher age gaps across multiple modules, whereas only two agents showed lower age gaps in more than one module (Figure S20 and Table S28). Notably, premarin (625 μg tablet) and glucosamine were associated with lower age gaps in several cellular systems, including endothelial, granulocyte-barrier, pancreas-epithelial, and liver-hepatocyte modules (Figures 6G and 6H).

Finally, metabolomic profiling revealed extensive metabolic correlates of cellular aging. Across 250 circulating metabolites, 1,780 of 3,500 metabolite-module associations were significant, with 237 metabolites (94.8%) associated with at least two cellular modules (Table S29). Lipoprotein and lipid metabolism represented the largest category of significant associations, with triglyceride-rich lipoproteins showing particularly widespread relationships across modules (Figure S21). Creatinine, a key marker of renal function, was significantly associated with all 14 cellular modules and ranked among the strongest associations in most modules (Figure 6I), whereas glycoprotein acetyls, a marker of systemic inflammation, were also broadly associated with cellular aging (Figure 6J).

Together, these findings identify both shared and module-specific correlates of cellular aging and demonstrate that distinct cellular aging programs are differentially associated with lifestyle, pharmacological, and metabolic factors.

## Discussion

In this study, we show that human cellular aging is organized into a modular architecture comprising conserved cross-organ and organ-specific cellular systems. By integrating a multi-organ single-cell transcriptomic atlas with population-scale plasma proteomics, we reveal that heterogeneous cellular aging modules are organized into conserved biological programs that differ in their temporal dynamics, functional remodeling, and clinical relevance. Module-specific aging states capture substantial inter-individual heterogeneity and are associated with disease burden, future disease risk, mortality, and modifiable health-related factors. Together, our findings provide a conceptual framework for understanding human aging as a coordinated yet modular cellular process and establish a foundation for precision geroscience.

A central finding of this study is that human cellular aging is organized into both conserved cross-organ and organ-specific modules. Most organ-cell type pairs clustered according to cellular identity rather than tissue of origin, indicating that related cell lineages share coordinated aging patterns despite residing in distinct anatomical environments^12,13,20^. This extends previous single-cell studies showing that lineage-associated molecular programs are largely preserved during aging^19,20,32,33^, and suggests that intrinsic cellular identity is a major organizing principle of human biological aging^34,35^. At the same time, highly specialized cell types, including hepatocytes, pancreatic epithelial cells, hematopoietic cells, and bone marrow stem cells, formed organ-specific modules, indicating that tissue-specific physiological functions further diversify aging in specialized cellular systems^12,36^. A small number of exceptions, such as muscle granulocytes and stomach endothelial cells, clustered outside their canonical lineages, suggesting that local tissue environments can occasionally reshape lineage-associated aging through shared microenvironmental influences^30,31^. For example, extracellular matrix remodeling, inflammatory signaling, and tissue repair processes are known to reshape cellular states during aging^14,37–39^. Together, these findings support a hierarchical organization of cellular aging in which lineage-dependent programs provide the primary framework, while tissue-specific functions further refine aging patterns.

Our results further demonstrate that cellular aging is temporally organized rather than synchronized across biological systems. DE-SWAN analysis revealed that structural and tissue-resident modules underwent molecular remodeling earlier in adulthood than immune modules, suggesting a sequential progression from tissue maintenance toward immune dysfunction^40^. This temporal ordering suggests that aging may initially compromise tissue architecture and metabolic homeostasis before progressively affecting immune competence^14,41–43^. Beyond differences in onset, cellular modules displayed four distinct aging patterns, including gradual accumulation, late-life acceleration, dynamic fluctuation, and young-state loss dominant. Structural modules generally exhibited gradual accumulation, consistent with the continuous decline of tissue maintenance and extracellular matrix homeostasis during adulthood^14^. In contrast, late-life acceleration observed in bone marrow stem cells and pancreatic epithelial cells may reflect age-dependent exhaustion in regenerative capacity and tissue physiological reserve^44,45^. The aging pattern of the B cell module further suggests greater responsiveness to environmental or systemic influences^46^. Notably, the erythroid module was characterized primarily by progressive depletion of youthful states rather than marked accumulation of accelerated aging, suggesting a potentially distinct aging pattern that warrants further investigation. These findings indicate that different biological systems age at different rates and through distinct trajectories, providing a potential explanation for the marked inter-individual heterogeneity observed in biological aging.

The clinical relevance of this modular architecture was supported by consistent associations with prevalent disease, incident disease, and mortality. Among the identified modules, epithelial, stromal, and myeloid aging showed particularly broad associations, underscoring the importance of barrier integrity, tissue homeostasis, and innate immune function in age-related diseases^47–49^. For example, accelerated epithelial aging may contribute to metabolic disease susceptibility, as epithelial dysfunction and impaired tissue repair have been implicated in insulin resistance and diabetes progression^50,51^. Individuals with accelerated aging across multiple modules exhibited progressively greater disease burden, incident disease risk, and mortality, suggesting that adverse outcomes reflect the cumulative disruption of multiple biological compartments rather than dysfunction of a single cellular lineage. Longitudinal analyses further demonstrated that the temporal relationship between aging and disease differs across modules, with some cellular modules exhibiting accelerated aging before clinical diagnosis, whereas others continuing to diverge following disease onset^52^.

Cellular aging also appeared to be influenced by modifiable factors beyond chronological age. Module-specific age gaps were associated with lifestyle, anthropometric, socioeconomic, and dietary factors, supporting the concept that biological aging reflects cumulative environmental and physiological exposures throughout life^53^. For example, adverse lifestyle factors, including smoking, sedentary behavior, and socioeconomic deprivation, were associated with accelerated aging across multiple cellular modules, whereas favorable behaviors and better physical fitness showed protective associations^54–57^. However, these associations were not uniform across cellular systems, suggesting that environmental exposures interact with cell-type-specific aging programs rather than affecting all biological systems uniformly. Similarly, circulating metabolic profiles highlighted lipid metabolism and inflammation as major correlates of cellular aging, with triglyceride-rich lipoproteins and glycoprotein acetyls showing widespread positive associations across modules, whereas HDL-related measures showed predominantly protective associations^42,58,59^. Likewise, the associations between several medications and lower age gaps in specific modules raise the possibility that different cellular systems may differ in their responsiveness to pharmacological interventions, providing a rationale for future cell-informed geroscience strategies^60–62^.

Several limitations should be acknowledged. First, although plasma proteins provide accessible readouts of systemic aging, circulating protein levels are influenced by secretion, turnover, and clearance, which may limit precise attribution to their tissue or cellular origins^63^. Second, although the identified cellular aging modules showed cross-organ coherence and functional consistency, they reflect patterns captured by currently available human single-cell reference atlases and may be refined as larger and more diverse cellular maps become available. Finally, the UK Biobank cohort predominantly comprises individuals of European ancestry, and validation in more diverse populations will be important to establish the generalizability of these findings.

In summary, we reveal a modular architecture of human cellular aging by bridging single-cell transcriptomics with population-scale plasma proteomics. Our findings show that cellular aging is organized into a modular architecture comprising conserved cross-organ and organ-specific aging modules with distinct temporal dynamics and clinical relevance. By providing scalable measures of coordinated cellular aging across biological systems, this framework offers a foundation for investigating aging mechanisms, understanding individual differences in aging trajectories, identifying potential cellular intervention targets, and advancing precision geroscience strategies.

## Methods

### Human multi-organ single-cell transcriptomic data

Single-cell transcriptomic data were obtained from the Tabula Sapiens 2.0 release^27^, a comprehensive human cell atlas generated by the Tabula Sapiens Consortium. The dataset comprises over 1.1 million single-cell RNA sequencing (scRNA-seq) profiles from 28 organs of 24 donors (11 males and 13 females; aged 22-74 years), encompassing 34 broad cell classes and approximately 300 annotated cell types. Tissue collection, processing, and dissociation were performed using standardized protocols across participating centers, as previously described^27,28^. Processed metadata were accessed through the Chan Zuckerberg CELL by GENE (CZ CELLxGENE) platform.

Quality control was applied to the cell-by-gene count matrix. Cells with fewer than 200 or more than 6,000 detected genes, or with >10% mitochondrial transcripts, were excluded. Genes detected in fewer than three cells were removed. Because circulating plasma proteins predominantly originate from vascularized tissues^23^, analyses were restricted to 16 organs with established contributions to protein secretion or leakage into the bloodstream, including blood, vasculature, bone marrow, spleen, heart, lung, liver, stomach, bladder, pancreas, kidney, large intestine, small intestine, muscle, skin, and adipose tissue. Broad cell types represented by fewer than 200 cells across the dataset were excluded owing to insufficient cell numbers for reliable downstream analyses. The final dataset comprised 691,744 cells from 20 donors (11 males and 9 females; aged 22-74 years), encompassing 22 broad cell types and 327 annotated cell types (Table S1). All analyses complied with the data usage policies of the Tabula Sapiens Consortium.

### Plasma proteomic data

Plasma proteomic data were obtained from the UK Biobank Pharma Proteomics Project (UKB-PPP)^29^, which profiled plasma samples from over 50,000 participants using the Olink Explore 3072 Proximity Extension Assay (PEA) platform. The platform quantified 2,923 unique proteins across eight panels. Proteomic measurements were available at up to three visits (UKB field ID 30900): baseline (instance 0; 2006-2010), first imaging visit (instance 2; from 2014), and repeat imaging visit (instance 3; from 2019). Normalized protein expression values obtained from the UKB were used for subsequent analyses.

Quality control excluded participants who had withdrawn consent (withdrawal list updated March 2026), proteins with >20% missing values, and individuals with >49.9% missing measurements. Remaining missing protein values were imputed using the K-nearest neighbors (KNN) algorithm implemented in the KNNImputer function of scikit-learn (version 1.7.2) with 10 nearest neighbors (k = 10)^64^. After quality control, the baseline dataset included 49,388 participants and 2,911 proteins. For longitudinal analyses, a separate cohort comprising individuals with measurements available at all three visits was constructed, including 991 participants and 1,460 consistently measured proteins. Participant and protein characteristics are summarized in Tables S2 and S3. All participants in the UK Biobank provided written informed consent at recruitment, and the present study was conducted under UK Biobank’s approved data access procedures^65^. Access to UK Biobank data was granted under application number 46387.

### Participants and phenotypes

Analyses were restricted to UK Biobank participants with available Olink Explore proteomics data. Phenotypic information was accessed through the UK Biobank data portal in August 2025, and follow-up was censored according to data-provider availability (https://biobank.ndph.ox.ac.uk/showcase/exinfo.cgi?src=Data_providers_and_dates). At baseline, participants were aged 39-70 years; 54% were female, and 93% were of White ethnicity. Maximum follow-up was approximately 17 years.

#### Clinical disease outcomes

Disease diagnoses were derived from hospital inpatient records (Field 41270), which contain ICD-10 codes recorded across all hospital episodes. Dates of first diagnosis were obtained from Field 41280, enabling reconstruction of longitudinal disease histories. The baseline assessment date (Field 3166), corresponding to blood collection for proteomic profiling, was used as the reference time point. Diseases diagnosed before baseline were classified as prevalent, whereas those diagnosed after baseline were considered incident. For prevalent analyses, individuals who developed the disease during follow-up were excluded to minimize outcome misclassification. For incident analyses, participants with a prior diagnosis of the disease of interest were excluded, and time-to-event analyses were performed from baseline until first diagnosis. Controls were defined as participants without the corresponding disease. Disease definitions were harmonized using established endpoints from previous studies and the FinnGen project (https://www.finngen.fi/en/researchers/clinical-endpoints), following their quality-control framework^66^. Only diseases with at least 1,000 cases were retained. This yielded 162 incident diseases, of which 53 were also included in prevalent disease analyses. Detailed disease definitions are provided in Tables S4 and S5.

#### Mortality data

Mortality information was obtained through linkage to national death registries. Data for England and Wales were provided by NHS England (censoring date: 31 August 2024), and data for Scotland by the NHS Central Register (censoring date: 30 November 2024). Records included date of death (Field 40000) and primary cause of death (Field 40001). Participants were followed from baseline until death or administrative censoring, whichever occurred first. During ∼17 years of follow-up, 5,411 deaths (11%) were recorded.

#### Health-related traits

A broad set of health-related traits was analyzed, including lifestyle factors (diet, physical activity, smoking, and alcohol use), socioeconomic and environmental variables, mental and cognitive health, anthropometric traits. For variables with repeated measurements at the same assessment visit, mean values were used. Responses such as “Do not know” or “Prefer not to answer” were treated as missing. To reduce potential confounding by disease status, individuals diagnosed with the corresponding disease before or at baseline were excluded from analyses of each trait. Phenotypes with fewer than 1,000 participants were excluded; binary traits additionally required at least 100 cases. A total of 144 traits were included, with sample sizes ranging from 1,216 to 18,712 participants (Table S6).

#### Medication use

Medication data were obtained from Field 20003, which captures self-reported regular medication use collected through verbal interviews. This field includes long-term prescribed medications but excludes short-term treatments, over-the-counter drugs, and dietary supplements. Information on vitamin and mineral supplements use was additionally obtained from Fields 6155 and 6179. Each medication or supplement was encoded as a binary variable (user vs. non-user), and only exposures reported by at least 100 participants were retained. In total, 151 exposures (138 drugs and 13 supplements) were included in downstream analyses (Table S7).

#### Plasma metabolic traits

Plasma metabolic profiling was generated using high-throughput nuclear magnetic resonance (NMR) spectroscopy on the Nightingale Health platform (UK Biobank Category 220). Detailed experimental protocols have been described in the UK Biobank NMR documentation (https://biobank.ndph.ox.ac.uk/showcase/ukb/docs/NMR_companion_phase2.pdf). A total of 250 NMR-derived metabolic and lipoprotein traits were quantified and grouped into nine biochemical categories: lipoprotein and lipids (192 traits), fatty acids (18), cholesterol (15), amino acids (10), glycolysis related metabolites (5), ketone bodies (4), apolipoproteins (3), fluid balance markers (2), and inflammation markers (1). To improve distributional properties and comparability across these traits, extreme values were excluded by removing observations exceeding four interquartile ranges from the median. Trait values were subsequently log-transformed and standardized to z-scores. To minimize potential confounding by pre-existing disease, individuals diagnosed with the disease before or at baseline were excluded prior to analysis.

### Identification of organ-cell type-enriched protein features

We used the Tabula Sapiens single-cell atlas to identify candidate plasma protein markers for organ-cell type pairs. Differential expression analysis was performed separately for each cell group against all remaining cells using the t-test implemented in the rank_genes_groups function of Scanpy python package (version 1.11.1)^67^. Statistical analysis was assessed using Student’s t-tests comparing each organ-cell type pair against all other cells, *P* values were adjusted for multiple testing using the Benjamini-Hochberg procedure. Genes were defined as differentially expressed if (i) false discovery rate (FDR) <0.05; (ii) log2-fold change (log2FC) ≥2; and (iii) mean expression ranked in the top 10% within the target cell type.

Differentially expressed genes were mapped to age-related plasma proteins measured in the UK Biobank. Associations between plasma protein and chronological age were evaluated using separate multivariable ordinary least-squares (OLS) linear regression models implemented in the statsmodels Python package (version 0.14.4), with adjustment for the predefined covariates. P values were adjusted for multiple testing using the Bonferroni correction, *P* value < 1.72 × 10⁻^5^ (0.05/2911) considered statistically significant. Analyses were restricted to vascularized organs with established plasma contributions, while reproductive organs were excluded to reduce sex-related confounding. This yielded 128 organ-cell pairs and 1,369 protein features across 16 organs and 22 broad cell types. Because some cell types were shared across organs, resulting features represent organ-cell-enriched signatures rather than strictly specific markers.

### Organ-cell type aging models and age gap estimation

Plasma proteomic data from 4,989 healthy individuals (40-70 years, 53.24±7.75) were randomly split into training (80%) and test (20%) sets. Biological age prediction models were developed using seven machine-learning algorithms, including LASSO regression, elastic-net regression, support vector regression (SVR), random forest regression, OLS linear regression, tree-based XGBoost and linear XGBoost, implemented using the scikit-learn and XGBoost (version 3.0.0) Python packages. Hyperparameters were optimized using grid search with fivefold cross-validation implemented in the GridSearchCV function of scikit-learn. The optimal model was selected according to the lowest mean squared error.

LASSO regression was selected based on consistent performance. Model robustness was assessed via 100 bootstrap iterations, with feature stability defined as the proportion of non-zero coefficients across runs. Model performance was evaluated in both training and test sets using mean squared error (MSE), mean absolute error (MAE), coefficient of determination (R2), and Pearson correlation coefficients (*r*). Predicted ages were calibrated using locally weighted scatterplot smoothing (LOWESS) implemented in the lowess function of the statsmodels package, followed by linear interpolation using interp1d from SciPy (version 1.15.3). Age gaps were calculated as predicted age minus LOWESS-estimated expected age and subsequently standardized to z scores within each model.

### Hierarchical clustering of organ-cell type aging models

Hierarchical clustering was used to characterize similarity among organ-cell type aging models based on their age gap profiles across individuals. Pairwise Pearson correlation coefficients were calculated between the age-gap values of all organ-cell type models using the pearsonr function in SciPy. Pairwise distance was defined as *d* = 1 - *r*, where *r* denotes the Pearson correlation coefficient between organ-cell type-level age gaps. Agglomerative hierarchical clustering was performed on the precomputed distance matrix using average linkage and a distance threshold of 0.55 with the AgglomerativeClustering function in scikit-learn. The clustering structure was visualized using a dendrogram generated in iTOL^68^, resulting in 14 major cellular aging modules. In addition, t-SNE was applied to the organ-cell type-level age gap matrix to provide a two-dimensional representation that further supported module interpretation.

### Construction of module-level aging state models

Module-level state models were built using the same framework applied to organ-cell type-level models. Differentially expressed genes were identified for each module by comparing module-associated cells with all other modules (FDR < 0.05, log2FC ≥ 2, top 10% expression) and mapped to plasma proteins measured in the UK Biobank. In contrast to organ-cell type-level features, which were permitted to overlap across organ-cell type pairs, module-level features showed substantially lower overlap because cells were uniquely assigned to modules and features were derived using module-versus-all comparisons. Across the 14 modules, 685 unique age-associated proteins were identified, yielding largely distinct module-specific protein signatures. Plasma proteomic data were split into training (80%) and test (20%) sets, and bootstrap-aggregated LASSO models (100 iterations) were trained with adjustment for sex and ethnicity. Module-level age gaps were calculated as predicted age minus LOWESS-estimated expected age and standardized subsequently within each module using z-score transformation.

### Sliding-window analysis of module-specific proteomic remodeling

Differential expression-sliding window analysis (DE-SWAN) was performed to characterize age-dependent molecular remodeling across cellular aging modules^69^. DE-SWAN identifies non-linear age-associated changes by comparing protein levels between individuals younger and older than a given center age within consecutive age windows. Module-specific protein features were analyzed using a 5-year sliding window with a 5-year step size across ages 40-70 years. At each center age, individuals younger and older than the center age within the corresponding window were compared, whereas individuals exactly at the center age were excluded. For each protein, differences between age groups were assessed using linear regression while adjusting for relevant covariates. P values from all protein-by-age comparisons were corrected using the Benjamini-Hochberg procedure to control the false discovery rate (FDR). Proteins with FDR-adjusted *q* < 0.05 were considered significant, and the number of significant proteins identified at each age window was used to quantify module-specific temporal molecular remodeling.

### Functional enrichment analysis

To characterize the biological functions represented by each cellular aging module, we first performed gene set enrichment analysis (GSEA) at the single-cell transcriptomic level^70^. Differential expression analysis was conducted by comparing cells belonging to a given module with all remaining cells. Genes were ranked according to the following metric:

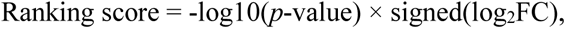

with log_2_FC used to break ties when ranking scores were identical. GSEA was performed using a weighted enrichment statistic and Gene Ontology Biological Process (GO-BP) annotations as the reference gene set database^71^. Gene sets containing 10-500 genes were retained for analysis. Normalized enrichment scores (NES) were calculated to quantify pathway enrichment, and statistical significance was determined by permutation testing with Benjamini-Hochberg correction (FDR < 0.05). Enrichment results were visualized using EnrichmentMap in Cytoscape^72^. Because module-specific marker genes were restricted to genes represented in the UK Biobank plasma proteomics dataset, enrichment analyses were performed using this matched gene universe as the background set rather than the whole genome to minimize bias arising from unmeasured genes.

To investigate molecular pathways associated with cellular aging in the population cohort, we further performed pathway enrichment analyses on plasma proteins associated with module-specific age gaps. For each aging module, proteins significantly associated with age-gap variation were identified using linear regression analyses. Proteins positively associated with age gap were classified as accelerated-aging proteins, whereas proteins negatively associated with age gap were classified as decelerated-aging proteins. The two protein groups were analyzed separately using over-representation analysis (ORA) implemented in g:Profiler web tool^73^. Statistical significance was evaluated using Fisher’s exact test and corrected for multiple comparisons using the Benjamini-Hochberg procedure.

### Association with clinical outcomes and traits

Associations between module-level age gaps and clinical outcomes were evaluated using regression-based models. Logistic regression was applied for prevalent diseases, whereas Cox proportional hazards models were used for incident diseases and all-cause mortality, with time since baseline as the underlying timescale. Associations between health-related traits, medication or supplement use, plasma metabolic biomarkers, and module-level age gaps were assessed using linear regression models, treating these variables as predictors and age gaps as outcomes.

All models were adjusted for relevant demographic, technical, lifestyle, and genetic covariates, as described below. Individuals with missing covariate data were excluded from the corresponding analyses. Effected estimates were reported as odds ratios (ORs), hazard ratios (HRs), or regression coefficients (β), together with 95% confidence intervals (CIs). Statistical significance was determined using Bonferroni correction with a threshold of *p* < 4.69 × 10^-6^, accounting for 761 tested phenotypes across 14 aging modules.

### Extreme aging stratification and clinical association analyses

Individuals with module-specific age-gap z scores ≥2 were classified as extreme accelerated agers, whereas those with z scores ≤-2 were classified as extreme decelerated agers. Participants with age-gap z scores between -2 and 2 were considered normally aging controls. Individuals exhibiting an extreme deviation (accelerated or decelerated) in at least one module were considered extreme aging individuals. Participants exhibiting an extreme deviation in exactly one module were classified as single-module extreme agers. Participants exhibiting extreme deviations in two or more modules, all in the same direction (either all accelerated or all decelerated), were classified as multi-module extreme agers and were further stratified according to the number of extreme modules (2, 3-4, or ≥5). Individuals simultaneously exhibiting accelerated aging in some modules and decelerated aging in others were classified as mixed agers and excluded from subsequent extreme-aging group analyses.

Associations between prevalent diseases and extreme aging groups were evaluated using logistic regression models. Logistic regression was also applied to incident diseases because many diseases had a limited number of incident events after extreme-aging stratification resulting in insufficient statistical power for Cox proportional hazards modeling. Associations between extreme aging groups and all-cause mortality were evaluated using Cox proportional hazards models. Associations were evaluated separately within each single-module extreme-aging group and across multi-module extreme-aging groups to assess whether disease risk increased with the number of extreme modules. All analyses were adjusted for the covariates.

For incident diseases, cumulative incidence functions were estimated using the cuminc function in the cmprsk R package (version 2.2.12) to compare incident disease risk in the presence of competing events (https://cran.r-project.org/web/packages/cmprsk/index.html), and group differences were assessed using Gray’s test. Follow-up time was defined as the interval between baseline proteomic assessment and disease diagnosis. Death was treated as a competing event. Participants who died before disease onset were censored at death, whereas those without the corresponding disease were censored at the UK Biobank administrative censoring date. Differences between aging groups were assessed using Gray’s test. For all-cause mortality, Kaplan-Meier survival curves were estimated using KaplanMeierFitter function, and differences between groups were assessed using the multivariate log-rank test from the lifelines (version 0.30.3) Python package^74^.

### Disease-centered longitudinal trajectory analysis

Longitudinal plasma proteomic data from UK Biobank participants with repeated measurements across instances 0, 2, and 3 were used to investigate temporal changes in module-specific aging surrounding disease onset. Module-specific age gaps at each visit were estimated using aging models trained in the cross-sectional cohort and applied to the longitudinal cohort. For each disease, repeated measurements from affected individuals were aligned relative to the time of diagnosis (time = 0), with negative and positive values indicating years before and after disease onset, respectively. Trajectories were generated only for cellular aging modules previously associated with each disease. For healthy controls, repeated measurements were aligned according to follow-up time relative to censoring. Temporal trajectories were visualized using locally weighted scatterplot smoothing (LOWESS), and 95% confidence intervals were estimated by bootstrap resampling.

### Statistical analysis

Unless otherwise specified, statistical analyses were performed using Python (version 3.12.10) and R (version 4.3.1). Associations of module-specific age gaps and extreme aging groups with clinical phenotypes were evaluated using regression models appropriate for the dependent variable. Continuous outcomes were analyzed using multivariable linear regression, binary outcomes using multivariable logistic regression, ordinal outcomes using ordinal logistic regression, and time-to-event outcomes using Cox proportional hazards regression, implemented using the ‘GLM’, ‘OrderedModel’, and ‘PHReg’ functions in the statsmodels Python package (version 0.14.4). Unless otherwise stated, regression models were adjusted for the predefined covariates, including sex (Field 31), ethnicity (Field 21000), Townsend deprivation index (Field 22189), fasting time (Field 74), season of assessment (Field 3166), body mass index (Field 21001), and the first ten genetic principal components (Field 22009) to account for demographic, lifestyle, and population structure-related confounding.

Multiple-testing correction was performed according to the analysis context. Differential expression and functional enrichment analyses were corrected using the Benjamini-Hochberg false discovery rate (FDR) procedure, whereas regression-based association analyses were corrected using the Bonferroni method according to the total number of statistical tests performed in each analysis. Unless otherwise specified, all statistical tests were two-sided. Hazard ratios (HRs), odds ratios (ORs), regression coefficients (β), and corresponding 95% confidence intervals (CIs) are reported where appropriate.

## Data availability

All model performance metrics and phenotype association results generated in this study will be publicly available upon publication. The datasets used in this study are publicly available. UK Biobank proteomics and phenotype data can be accessed by approved researchers through application to the UK Biobank resource (https://biobank.ndph.ox.ac.uk/). The data used in this study were obtained under application number 46387. Human multi-organ single-cell transcriptomic data were obtained from Tabula Sapiens (https://tabula-sapiens.sf.czbiohub.org/), and processed datasets are publicly available through the CZ CELLxGENE platform (https://cellxgene.cziscience.com/collections/e5f58829-1a66-40b5-a624-9046778e74f5).

## Code availability

Analyses code supporting this study have been deposited in the GitHub repository but is not publicly accessible in the current preprint version. Please contact us via email should you have any questions. Any additional information required to reanalyze the data reported in this paper is available from the lead contact upon request.

## Acknowledgements

This study used the UK Biobank Resource under application numbers 46387. We want to thank all the participants and researchers from the UK Biobank. This work was supported by the Science and Technology Projects of Xizang Autonomous Region (XZ202502ZY0036 to T.-L.Y.), Science Fund for Distinguished Young Scholars of Shaanxi Province (2025JC-JCQN-054 to Y.G.), the National Natural Science Foundation of China (32470639 to T.-L.Y., and 82401762 to J.G.), and China Postdoctoral Science Foundation (2024M762573 and 2026T190604 to J.G.). This work was also supported by the High-Performance Computing Platform and Instrument Analysis Center of Xi’an Jiaotong University.

## Author contributions

Conceptualization, T.-L.Y. and Y.G.; methodology, J.G. and C.-C.L.; investigation, J.G., C.-C.L. Y.X., F.J., J.-H.W., W.S., X.-L.Y., and H. D.; UK Biobank data acquisition and processing, T.-L.Y. and J.G.; writing – original draft, J.G.; writing – review & editing, T.-L.Y., Y.G., J.G., and C.-C.L.; visualization, J.G., C.-C.L.,Y.X., F.J. W.-J.H. and W.S.; constructive suggestions: S.-S.D., supervision, T.-L.Y. and Y.G.; funding acquisition, T.-L.Y., Y.G., and J.G..

